# CpG Atlas: A centralized multi-layer database and AI interface for DNA methylation research

**DOI:** 10.64898/2026.05.30.729020

**Authors:** Jenel F. Armstrong, Shawn Wahi, Daniel Borrus, Raghav Sehgal, Syed Rizvi, Shiyang Zhang, Macsue Jacques, Nir Eynon, David van Dijk, Albert Higgins-Chen

## Abstract

DNA methylation research has vastly expanded over the past decade, producing a wealth of epigenome-wide association studies, biomarker algorithms such as epigenetic clocks, technical performance analyses, and functional annotations for CpG sites. However, these resources remain fragmented across dozens of databases and supplementary files within manuscripts, forcing researchers to spend time and effort on data cleaning and integration prior to meaningful analyses. No single resource currently unifies this information into a centralized, easy-to-query framework. Here, we present CpG Atlas, a curated relational database that integrates 18 distinct annotation layers encompassing over 1.2 million CpG sites across all four generations of Illumina methylation arrays (HM450K, EPIC v1, EPIC v2, and MSA). Built on a snowflake schema with a canonical probe identifier hub implemented in SQL, CpG Atlas consolidates over 800,000 CpG-trait associations, results from Mendelian randomization analyses, CpG membership across 81 epigenetic clocks, array manifest information, and probe reliability data. It further includes specialized layers such as solo-WCGW, CoRSIVs, PRC2 binding, transposon and retroelement annotations, tissue-specific differentially methylated positions across 17 tissues, and hallmarks of aging and cancer. To maximize utility and ease of use, the database is paired with an interactive web tool and a natural language-to-SQL query interface, enabling users to quickly perform complex multi-dimensional queries. Detailed documentation about every data source and table is also provided, facilitating the identification and interpretation of relevant studies. We demonstrate the utility of CpG Atlas through two case studies: a systematic enrichment analysis revealing distinct functional signatures across 16 epigenetic clocks, and an iterative biomarker discovery workflow for IBD that leverages cross-layer integration. Because it is readily scalable simply by adding or updating tables in the database, CpG Atlas provides a continuously evolving and extensible infrastructure for the epigenetics community that supports collaborative research, interpretable biomarker development, and integrative analyses across the growing landscape of epigenetic data.

## Introduction

DNA methylation (DNAm) at cytosine-guanine dinucleotides (CpG sites) represents one of the most commonly and comprehensively assayed epigenetic modifications to the human genome. The widespread adoption of Illumina’s Infinium bead array technology, including the HumanMethylation450 (450K), MethylationEPIC v1.0 (EPIC), MethylationEPIC v2.0 (EPICv2), and Methylation Screening Array (MSA) platforms, has enabled cost-effective measurement of DNAm at hundreds of thousands of genomic loci across hundreds of thousands of human samples (1–4). The great amount of methylation data has set the stage for numerous Epigenome-Wide Association Studies (EWAS) linking CpG sites to diseases and outcomes (5), dozens of validated epigenetic clocks predicting chronological age, biological age, and tissue-specific aging (6, 7), and an expanding catalog of DNAm-based biomarkers for proteins, metabolites, disease risk, and therapeutic response (8–10). DNAm has proven particularly powerful in aging research: epigenetic clocks based on small panels of CpGs can predict chronological age with remarkable precision (11, 12) and “second-generation” clocks trained on mortality outcomes outperform many conventional risk factors in predicting healthspan and lifespan (13, 14).

Despite these remarkable advancements, there remains a gap in translating DNA methylation knowledge into robust, reproducible, and biologically meaningful biomarkers. This gap affects two major groups of researchers: those interpreting EWAS results, and those building new DNAm biomarkers. For researchers interpreting lists of EWAS hits, establishing clear mechanistic insights is critical for downstream applications. For example, when annotating novel EWAS hits associated with traits or diseases, researchers typically must perform tedious manual cross-referencing by querying their resulting CpG lists against EWAS databases to identify prior associations, followed by entirely separate functional enrichment analyses using disconnected tools to derive biological meaning (15, 16). For researchers building novel biomarkers, feature selection is a critical step, such as pre-selecting CpGs with adequate test-retest reliability (17, 18) or pre-selecting specific subsets of CpGs based on targeted biological annotations to build predictive models, as seen in the development of the causality-enriched clocks, RetroAge, and PathwayAge (19–21). However, both workflows currently rely on labor-intensive, manual harmonization of data. Researchers are typically forced to annotate or filter their CpG lists using only one or two functional layers at a time, leaving fundamental questions about how these diverse annotations interact entirely unaddressed. Furthermore, the requisite computational skills also exclude many clinicians, biologists, and epidemiologists who might be best positioned to translate these findings.

Existing resources have made important contributions to solving this CpG annotation problem, but remain incomplete for the holistic task of centralizing all the available CpG annotations into one place, as they often focus on a limited number of CpG annotation layers. The EWAS Open Platform and EWAS Catalog (22, 23) provide invaluable repositories of curated EWAS associations and causal relationships. KnowYourCG (24) provides elegant CpG-level functional screening against diverse biological knowledge bases. Packages like methylCIPHER and Biolearn provide CpG contributions to various aging biomarker algorithms (25, 26). However, there is not yet any resource that addresses the fundamentally heterogeneous nature of CpG probes: a single CpG identified in an EWAS may simultaneously represent an aging hallmark, a poorly reproducible measurement, a tissue-specific epigenetic landmark, a clock component, and a target of a major transcription factor. Currently, no existing database bridges these distinct domains to provide a unified platform for CpG list contextualization and iterative, quality-controlled biomarker design.

Fundamentally, the field faces two major bottlenecks: the fragmentation of DNAm information, which prevents broad accessibility for those without technical expertise, and the continuous influx of new annotations, which demands an infinitely scalable infrastructure. To address these challenges, we present CpG Atlas, a comprehensive knowledgebase and AI-powered query interface. CpG Atlas centralizes 18 curated functional data layers into a single, scalable database schema. To democratize access to this complex data, the database is paired with a browser-based interface and a natural language-to-SQL query system. Supported by detailed wiki documentation, this architecture allows users to execute highly complex, multi-layered queries using plain English, bypassing the tedious work of manual data curation and harmonization. We demonstrate the utility of CpG Atlas through two applications: a systematic enrichment analysis of existing epigenetic clocks and an iterative, quality-controlled discovery of highly reliable CpGs for an inflammatory bowel disease epigenetic biomarker.

## Results

### The CpG Atlas database: 18 integrated functional annotation layers

CpG Atlas is built upon a DuckDB (27) relational database hosting 18 harmonized data tables, covering data from over 20 peer-reviewed publications (Figure 1a, Supplementary Table S1). We organized these layers into six functional categories based on their primary utility in the biomarker design and analysis workflows: (1) Array Manifests and Technical Quality, (2) Causal and Associational Evidence, (3) Functional and Biological Context, (4) Tissue Specificity, (5) Repetitive Elements, and (6) Clock Membership.

**Figure 1.**
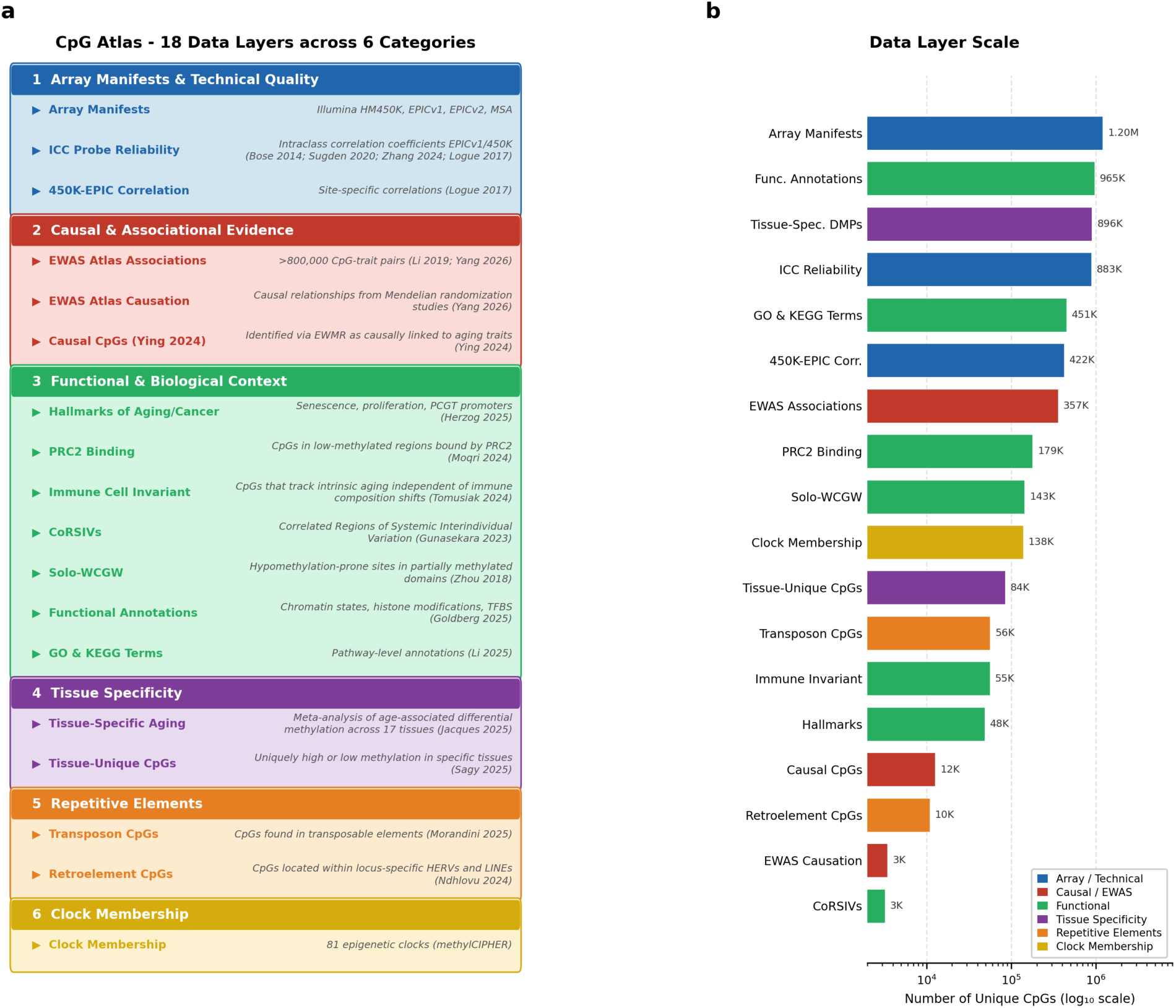
The CpG Atlas Data Ecosystem. **a.** Overview of the 18 integrated functional layers organized into six categories: 1) Array Manifests and Technical Quality (array manifests for 450K/EPICv1/EPICv2/MSA and ICC reliability from 4 studies), 2) Causal and Associational Evidence (EWAS Atlas associations, EWAS causation, and causal CpGs from EWMR), 3) Functional and Biological Context (Hallmarks of Aging/Cancer, PRC2 binding, immune invariant CpGs, CoRSIVs, Solo-WCGWs, functional annotations, GO/KEGG pathways), 4) Tissue Specificity (tissue-specific aging DMPs and tissue-unique CpGs), 5) Repetitive Elements (transposon CpGs, and retroelement CpGs) and 6) Clock Membership in 81 epigenetic clocks. **b.** Horizontal bar chart showing the number of unique CpGs covered by each data layer on a log₁₀ scale, color-coded by category.

#### (1) Array Manifests and Technical Quality

CpG Atlas incorporates complete array manifest files for all four major Illumina methylation array platforms: the HumanMethylation450 BeadChip (450K; 485,553 loci; Build hg19 (28)), MethylationEPIC v1.0 (EPICv1; 865,918 loci; Build hg19 (29)), MethylationEPIC v2.0 (EPICv2; 937,055 loci; Build hg38 (30)), and the Methylation Screening Array (MSA; 281,806 loci; Build hg38 (31)). The manifests provide genomic coordinates, Infinium probe design type (I or II), and detailed genetic context including gene associations from RefSeq and GENCODE, CpG island relationships (Island, Shore, Shelf), regulatory features such as enhancers and DNase hypersensitivity sites, and SNP annotations derived from dbSNP. CpG Atlas explicitly cross-references probe identifiers across all four arrays, enabling seamless conversion and comparison across platforms, which is essential given that most of the older DNAm datasets use the 450K or EPICv1 arrays while newer cohorts are starting to generate EPICv2 or MSA data.

Another defining feature of CpG Atlas is its integration of probe technical reliability data, which is essential for reproducibility, yet is inconsistently integrated into DNA methylation analyses. DNAm measurements from Illumina arrays show highly variable test-retest reliability across probes, with intraclass correlation coefficients (ICC) ranging from near zero to near perfect across the same array platform (32). CpG Atlas integrates ICC values from four independent studies totaling 764 replicate sample pairs: Bose et al. (n=265 replicates, ARIC study, 450K (33)), Sugden et al. (n=350, Dunedin study, 450K (32)), Zhang et al. (n=138, ADNI study, EPICv1 (34)), and Logue et al. (n=11, TRACTS study, EPICv1 (35)). In total, the ICC datasets cover over 880K unique CpG sites. Additionally, site-specific 450K-EPIC cross-platform Pearson correlations from Logue et al. (n=145) for 422,524 shared probes further enable researchers to assess whether their findings generalize across array generations (35).

#### (2) Causal and Associational Evidence

CpG Atlas sources EWAS associations from the EWAS Open Platform (22, 36, 37), which include over 800,000 manually curated CpG-trait associations from 1,568 publications covering 880 distinct traits spanning diseases, environmental factors, behaviors, and clinical biomarkers. This provides immediate phenotypic context for a CpG of interest. Beyond statistical associations, CpG Atlas integrates 2 levels of causal evidence for CpG-trait relationships: the EWAS Causation module from Yang et al (2026) (22), and the causal CpG catalog from Ying et al. (2024) (19). The EWAS Causation table integrates 17,494 causal relationships derived from published Mendelian randomization studies, including 3,836 methylation→trait, 45 trait→methylation, and 10,557 methylation→gene expression causal links. The causal CpGs table from Ying et al. (2024) includes CpG sites identified via epigenome-wide Mendelian randomization (EWMR) to be causal to 8 aging-related traits. The causal and associational evidence tables in CpG Atlas preserve all MR effect estimates, EWAS statistics, and study information, enabling users to apply their own evidence thresholds and study choices when filtering for associational or causal CpG.

#### (3) Functional and Biological Context

A major layer in CpG Atlas is the integration of mechanistic and biological annotations beyond genomic coordinates to explain how CpG methylation changes with age and disease. Seven complementary tables address this matter. **1)** The **Hallmarks of Aging and Cancer** module from Herzog et al. (2025) (38) classifies CpGs into 3 mechanistically distinct categories: senescence-associated CpGs (8,399 sites altered in senescent cells across multiple senescence-induced and control cells datasets), proliferation-associated CpGs (39,221 sites whose methylation changes with proliferative status, derived from cells treated with proliferation inhibitors), and Polycomb Group Target (PCGT) promoter CpGs (2,577 sites within 200 bp of TSS of PCGT genes, unmethylated in fetal tissues). These categories in CpG-Atlas allow researchers to quickly pinpoint whether observed DNA methylation changes are driven by replicative exhaustion, developmental pathways, or oncogenic processes. **2)** The **PRC2 binding data** from Moqri et al. (2024) (39) provides continuous binding intensity scores for CpGs located within low-methylated regions (LMRs) in human embryonic stem cells, based on EZH2 and SUZ12 ChIP-Seq data from ENCODE in H1 hESCs. Moqri et al. show that PRC2-bound LMRs account for approximately 90% of age-dependent hypermethylation genome-wide in somatic cells, making this annotation crucial for identifying CpGs whose methylation gain during aging reflects a specific mechanistic process rather than stochastic changes. **3)** The **Immune Cell Invariant** module from Tomusiak et al. (2024) (40) addresses a critical confounder in blood-based DNAm data, changes in immune cell composition. Tomusiak et al. demonstrated that human naïve CD8+ T cells show an epigenetic age 15-20 years younger than effector memory CD8+ T cells from the same individual, and that many existing clock CpGs may reflect immune composition changes rather than pure cell-intrinsic aging. The 55,895 CpGs in the immune cell invariant table were selected by requiring a correlation with chronological age (|r| > 0.3) but no correlation with a sample being a naive CD8+ sample (|r| < 0.3). **4) CoRSIVs** (Correlated Regions of Systemic Interindividual Variation; Gunasekara et al., 2023 (41)) are regions where DNAm is highly variable between people but consistent across the different tissues of each person, functioning as epigenetic polymorphisms with exceptional mQTL effects (median R^2^ = 0.76). CpG Atlas maps 3,781 EPIC Array probes overlapping known systemic interindividual variation (SIV) regions, enabling researchers to identify whether their CpGs of interest represent tissue-consistent epigenetic variation. **5)** The **Solo-WCGW** annotation from Zhou et al. (2018) (42) identifies CpGs with the WCGW tetranucleotide motif (W = A or T) and no neighboring CpGs within +/-35 bp, located within Partially Methylated Domains (PMDs). These sites exhibit significant methylation loss with cell division across development and aging, functioning as a mitotic clock that correlates with chronological age, proliferation markers, and somatic mutations in cancer. The Solo-WCGW annotation helps researchers distinguish mitotic-driven methylation changes from those driven by transcriptional or environmental factors. **6)** The **Functional Annotations** module was derived from the KnowYourCG R package in R/Bioconductor (24). This table provides CpG-level annotations for transcription factor binding sites from ENCODE ChIP-seq consensus data, chromatin states, and histone modifications for 450K, EPICv1, and EPICv2 arrays, enabling access to regulatory context rather than mere genomic coordinate-based information. **7)** The **GO and KEGG** pathway annotations from Li et al. (2025) (21) map each CpG to Gene Ontology biological process terms and KEGG pathways. For each gene, the authors analyzed CpG sites located within a +/-20 kb region to capture both promoter-proximal and distal regulatory elements covering 2,779 distinct pathways from GO and 300 pathways from KEGG. This annotation creates a direct bridge between individual CpG associations and pathway-level biological interpretation.

#### (4) Tissue Specificity

CpG Atlas integrates two complementary resources for tissue-specific methylation context. The Tissue-Specific Aging DMP module from Jacques et al. (2025) (43) provides meta-analyzed age-associated differentially methylated positions (DMPs) across 17 tissues (adipose, blood, brain, breast, buccal, cervix, colon, heart, kidney, liver, lung, muscle, pancreas, rectum, retina, and skin) from 131 datasets totaling over 15,000 samples. DMPs were identified by multivariate linear regression adjusted for sex, BMI, and batch effects, combined across datasets by meta-analysis. DMP records include effect size, standard error, heterogeneity index (I2), and direction of change, enabling users to assess whether a candidate CpG is a conserved pan-tissue aging marker or a tissue-specific signal. Tissue-Unique CpGs module from Sagy et al. (2025) (44) identified 87,922 CpG sites across 22 tissues, derived from 5,323 samples by selecting sites where a tissue’s median methylation was at least 0.1 below the 5th percentile (uniquely low) or above the 95th percentile (uniquely high) of all other tissues. Together, these two tissue modules enable CpG Atlas users to answer important questions for interpreting aging biomarkers in accessible blood samples, like “Does this site change with age across tissues?” and “Is this site associated with loss (or gain) of tissue-specific epigenetic identity?”

#### (5) Repetitive Elements

Two layers address the biology of repetitive DNA elements: the Transposon CpG module (45) and the Retroelement CpG module (20). The Transposon CpG table uses RepeatMasker to annotate CpGs across all major transposable element classes: LINEs, SINEs, LTRs, and DNA transposons, at the family level, capturing per-probe methylation drift rates and youthful baseline methylation levels (at age 20). A residuals column further distinguishes active transposon de-repression from passive replication-driven hypomethylation. On the other hand, the Retroelement CpGs table provides locus-specific annotations of 60 HERV families and full-length LINE-1 elements mapped to EPIC probes using the Telescope pipeline. Together, these tables enable queries ranging from whether a given probe resides within a transposable element and how its methylation drifts with age, to identifying the precise HERV locus or retrotransposition-competent LINE-1 underlying a probe’s annotation.

#### (6) Clock Membership

The Clock Membership module in CpG Atlas provides a binary matrix indicating whether a given CpG belongs to any of the 81 published epigenetic clocks we extracted from the methylCIPHER package by accessing each clock’s component CpGs (25). This table integrates clocks predicting chronological age, biological age, mortality, pace of aging, clinical biomarkers, and cellular proportions from publications spanning 2013-2026 (Supplementary Table S2). This enables instant identification of probes that are widely used across clock generations, potentially revealing over-represented sites that may drive shared (or different) behavior among clocks.

### The CpG Atlas Explorer: a browser-based query and visualization interface

To make the database accessible without any programming expertise, we developed the CpG Atlas Explorer, a browser application that runs DuckDB-WebAssembly (DuckDB-WASM) (46) entirely client-side, with no server infrastructure required. All 30 database tables are exported to Apache Parquet format and hosted on Cloudflare R2 object storage, where the browser streams and queries them on demand. The Explorer is publicly accessible at *<u>cpg-atlas-pi.vercel.app</u>*. The interface is organized into five functional modules (Figure 2a). The **Overview** page provides access to every table with a summary; users may also upload their own tabular data (CSV or Excel) and later use it like an Atlas table. The **CpG Lookup** module enables instant retrieval of all annotation layers for one or more probe identifiers, presenting results as a joined table. The **SQL Query** module offers an easy-to-use and accurate NL-to-SQL query editor, and Plotly-based interactive visualization of results. The **CpG List Analysis** module accepts one (or multiple) user-supplied CpG list and automatically computes enrichment across selected annotation layers using Fisher’s exact test. The **Wiki** module provides embedded documentation for all 18 data layers, including method summaries, key statistics, and primary literature references, accessible within the tool. Rather than enforcing a linear analysis pipeline, these modules are designed to support an iterative, hypothesis-driven workflow. As illustrated in Figure 2b, researchers can smoothly navigate between different tabs to continuously refine their analyses based on evolving annotation results. For example, a user might begin with a biological annotation of interest, navigate to the SQL Query module, utilizing the AI assistant to extract an initial pool of candidate CpGs associated with a specific trait. By transitioning those results into the CpG List Analysis module, the researcher can instantly evaluate the list’s broader context. As the tool surfaces new biological insights, such as an unexpected enrichment for specific genomic regions, the researcher can then read more about this specific annotation in the associated Wiki page and refine their hypothesis accordingly. They can then return directly to the SQL Query or CpG Lookup tabs to execute more complex, narrowly tailored searches based on those fresh insights. This dynamic cycle of query, evaluation, and refinement empowers users to conduct deep exploratory data analysis.

**Figure 2.**
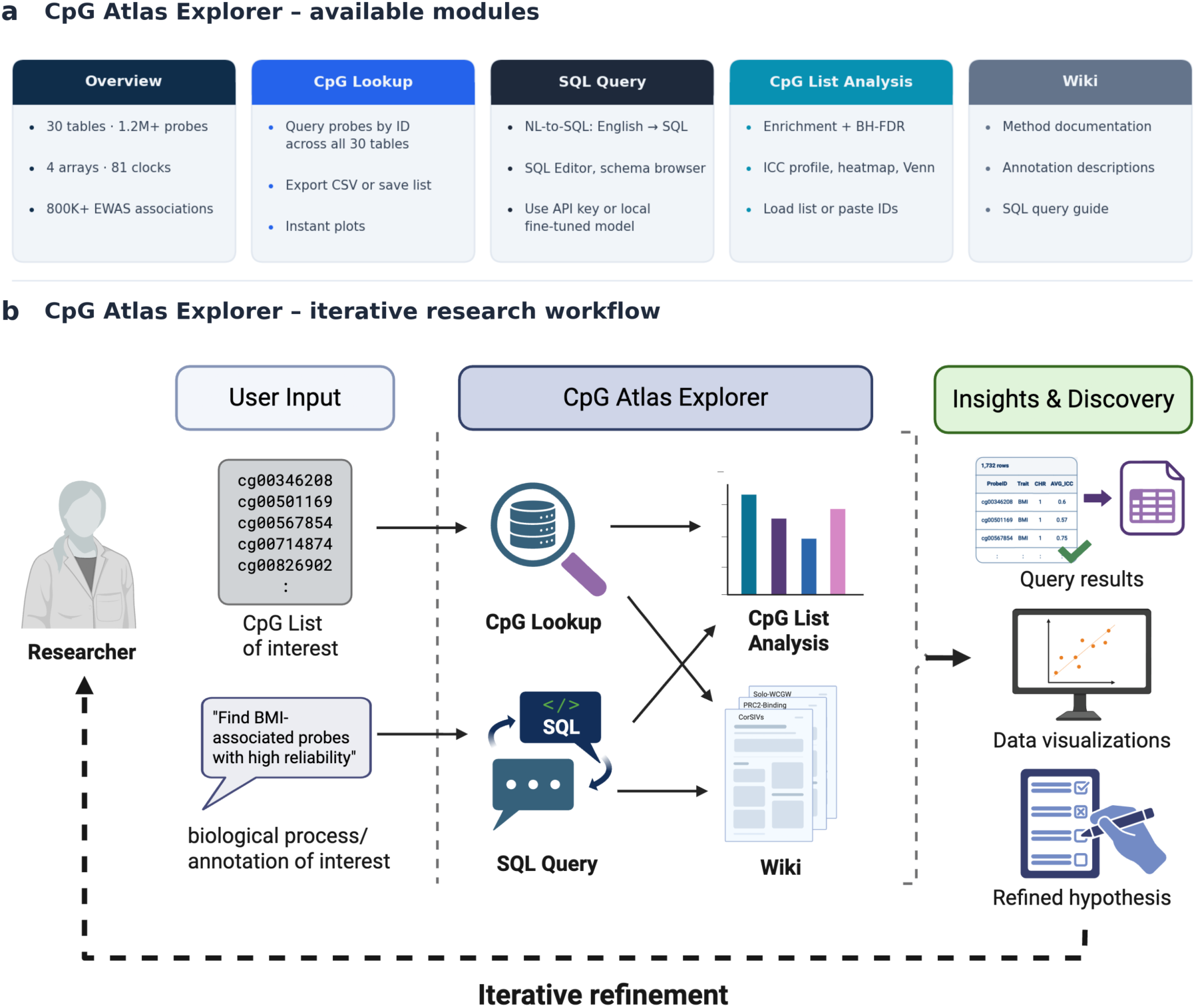
CpG Atlas Explorer interface and analytical workflows. **a.** Overview of the five functional modules: Overview, CpG Lookup, SQL Query (with NL-to-SQL), CpG List Analysis, and Wiki. **b.** Iterative research workflow.

### NL-to-SQL - A fine-tuned natural language to SQL model

While the Explorer’s CpG List Analysis and CpG Lookup modules cover common analytical workflows, formulating novel multi-table queries still requires SQL proficiency. To remove this barrier entirely for users, we developed a natural language-to-SQL (NL-to-SQL) query system integrated into the Explorer’s SQL Query tab (Figure 3). This allows researchers to describe their analytical intent in plain language, for example, “find all CpGs associated with mortality that have ICC greater than 0.75 and are immune cell invariant”, and receive an executable SQL query in return.

**Figure 3.**
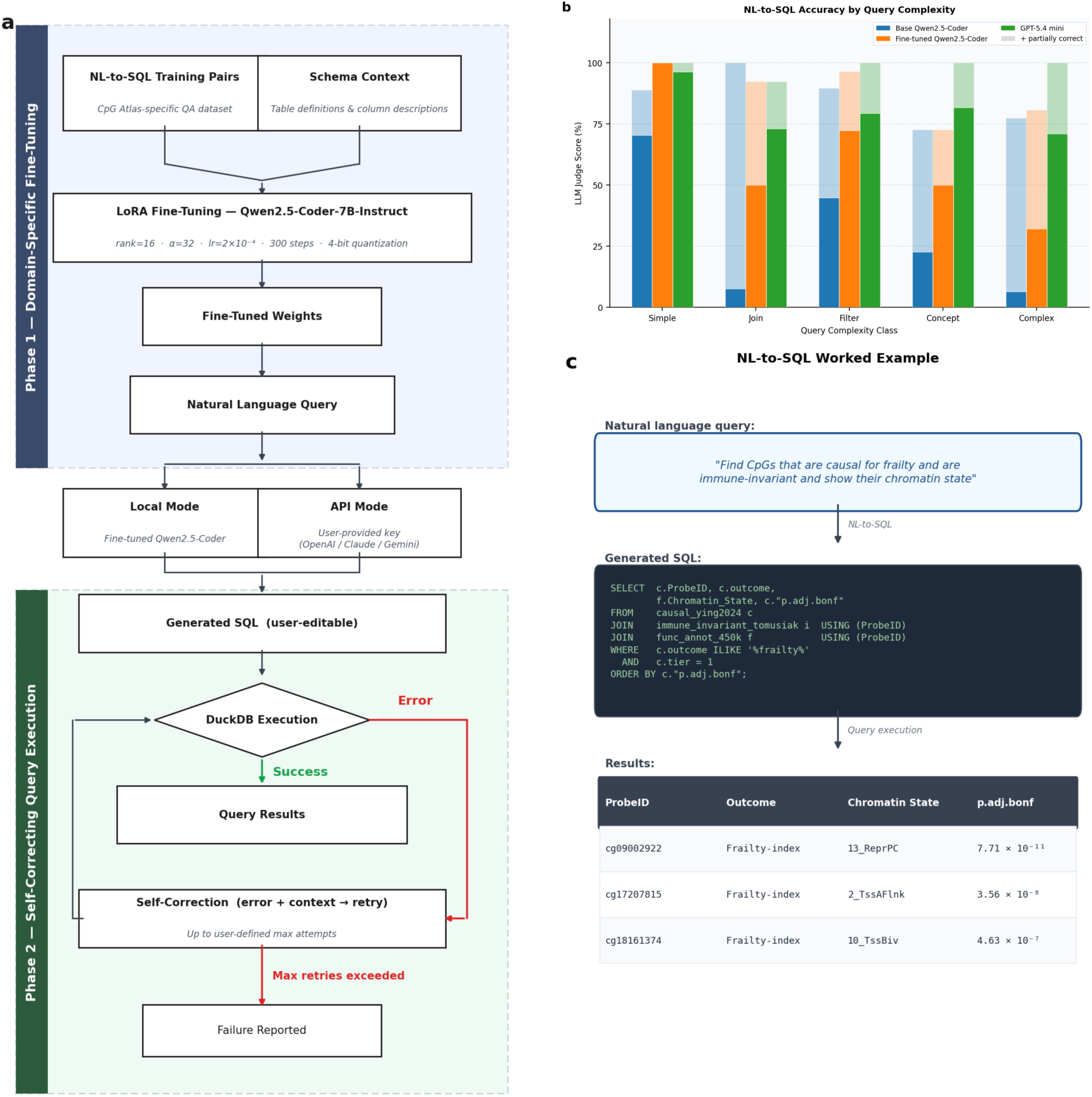
NL-to-SQL Model. **a.** Architecture of the NL-to-SQL system. Phase 1 involves fine-tuning of Qwen2.5-Coder-7B-Instruct on a CpG Atlas-specific training QCAs using LoRA. Users may choose between Local Mode, in which the fine-tuned model runs on their own hardware, or API Mode, in which an external provider (OpenAI, Anthropic, or Google Gemini) is queried using a user-supplied key. Phase 2 implements a self-correcting execution loop: the generated SQL is submitted to DuckDB; if an error is returned, the error message is fed back to the model for up to user-defined number of correction attempts before reporting failure. **b.** Query accuracy across five complexity categories evaluated on a held-out benchmark. **c.** Worked example illustrating a multi-table NL-to-SQL query.

The system operates in two phases (Figure 3a). Phase 1 is domain-specific fine-tuning: Qwen2.5-Coder-7B-Instruct (47) was fine-tuned using Low-Rank Adaptation (LoRA) (48). Phase 2 is self-correcting query execution: the generated SQL is passed to DuckDB for execution; if execution fails, the error message is appended to the prompt and the model is asked to revise its output, with the cycle repeating up to a user-defined number of attempts before a failure is reported. Researchers can run the system in two modes. In **Local Mode**, the fine-tuned Qwen model runs entirely on the user’s own hardware; no API key is required, and no data leaves the local environment, which is important for researchers working with sensitive or patient-derived methylation data under institutional data governance restrictions. In **API Mode**, users supply their own API key for a model of their choice (OpenAI GPT, Anthropic Claude, Google Gemini, etc.), which is then supplied with the database schema to handle query generation. Both modes produce a generated SQL query that the user can inspect and edit before execution. Across a held-out benchmark of 135 queries spanning five complexity classes: (1) simple single-table lookups, (2) multi-table joins, (3) filtering, (4) concept-mapping queries requiring biological synonym understanding, and (5) complex queries requiring an additional analysis step, the fine-tuned Qwen model achieves 61% overall accuracy with perfectly correct answers, compared to 30% using base model, and 88% overall accuracy with added partially correct answers, compared to 86% using base model (Figure 3b). We also tested the accuracy of the OpenAI GPT-5.4 mini model, which achieved 80% overall accuracy and 98% accuracy when partially correct answers are included (Figure 3b). Full evaluation metrics can be found in Supplementary Table S3.

### Systematic enrichment analysis reveals distinct functional signatures across epigenetic clocks

To illustrate the unique analytical capacity of a unified multi-layer database, we performed a systematic enrichment analysis of 16 major epigenetic clocks against 21 CpG Atlas annotation layers (Figure 4a, Supplementary Table S4). Assembling this analysis from scratch, which includes combining clock membership, ICC reliability, EWAS associations, causal evidence, hallmarks of aging, PRC2 binding, immune invariance, CoRSIV overlap, Solo-WCGW status, transposon and retroelement annotations, and functional genomic context, would require downloading and harmonizing data from more than a dozen independent sources. With CpG Atlas, it required a single database query. Statistical evaluation (using a Yates-corrected chi-squared approximation of Fisher’s exact test, followed by Benjamini-Hochberg FDR correction across all 336 tests) identified 143 significant associations (FDR < 0.05), demonstrating that the annotation layers in CpG Atlas carry substantial, non-redundant biological signal across different clocks.

**Figure 4.**
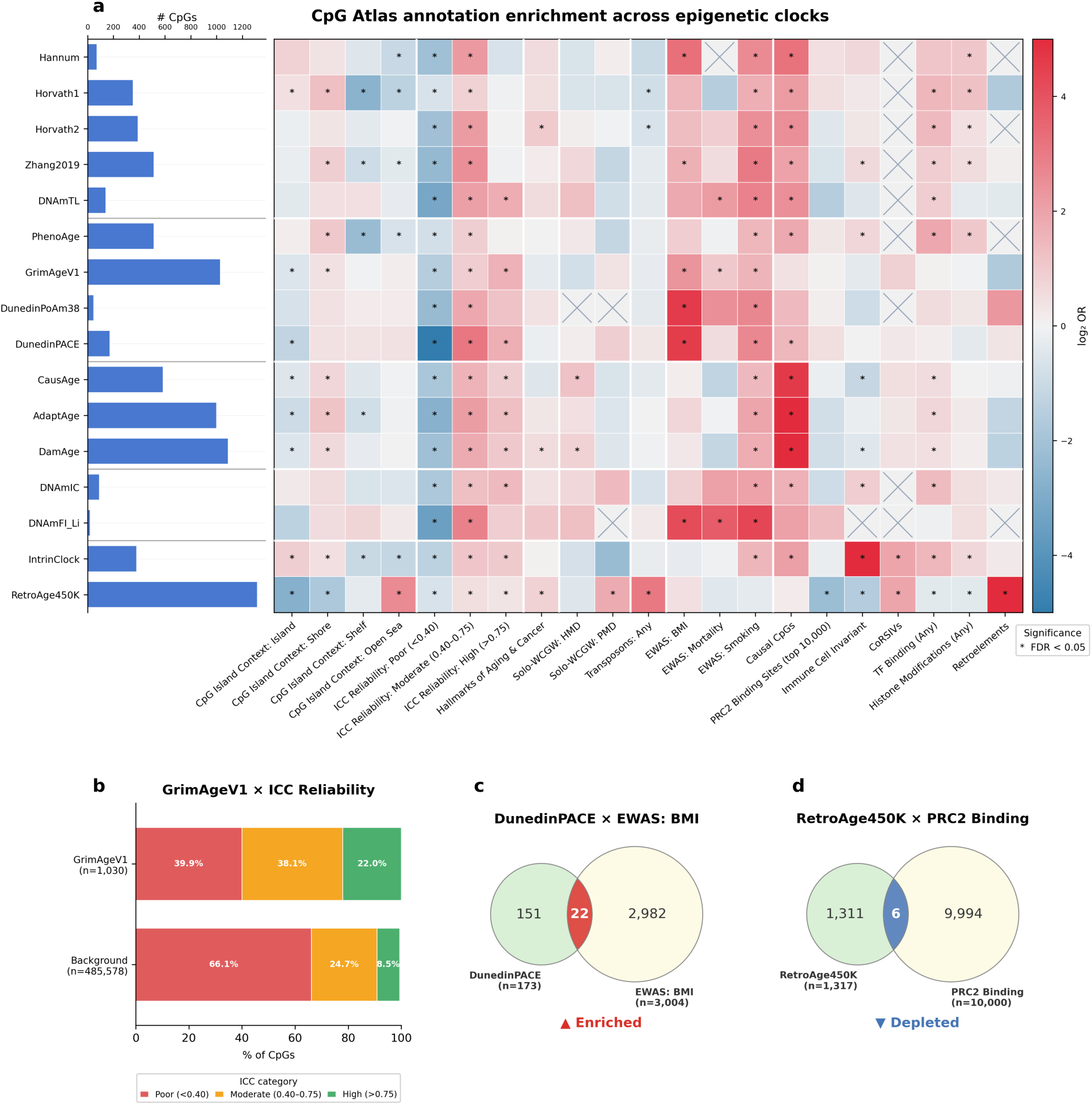
Clock Enrichment Analysis. **a**. Enrichment analysis heatmap for 16 clocks x 21 annotations. For each clock and annotation pair, a 2×2 contingency table is constructed comparing probe membership in the clock against the full Illumina 450K array background (N = 485,512 probes). Yates-corrected chi-squared approximation of Fisher’s exact test is applied and p-values are corrected using the Benjamini-Hochberg procedure across all 336 tests. Enrichment heatmap displaying log₂ odds ratios for all 16 clocks × 21 annotation combinations. Cells with asterisks are significant at FDR < 0.05 (143 of 336 tests). **b-d.** Selected annotation profiles highlighting clock-specific enrichment signatures.

### Expected Enrichments by Construction

Several results act as critical sanity checks, behaving exactly as expected by construction. RetroAge450K, defined entirely from retroelement probes, shows complete overlap with the Retroelements annotation (OR = ∞, FDR = 0). IntrinClock, built to track cell-intrinsic aging, shows complete overlap with the Immune Cell Invariant layer. The causal clocks (CausAge, DamAge, and AdaptAge) show the strongest enrichment for the Causal CpGs annotation (log_2_ *OR* = 4.62 to 5.65, FDR = 0), expected given all three were trained on the Ying et al. Mendelian randomization catalog. Additionally, smoking-associated CpGs are significantly enriched in 15 of the 16 evaluated clocks, reflecting the well-established, systemic impact of smoking across the methylome.

### Technical Reliability and EWAS Pre-selection

A critical pattern emerges when evaluating technical reliability (ICC). First-generation chronological age clocks, where the only pre-selection criterion was an association with age, show no enrichment for high-reliability CpGs. PhenoAge similarly lacks high-ICC enrichment, likely because it was trained to predict a composite outcome based on 9 biomarkers along with chronological age. This is consistent with our prior findings (17). In contrast, clocks with CpGs pre-selected on characteristics other than chronological age are highly enriched for reliable probes. GrimAge, for example, is a composite mortality predictor trained on plasma protein and smoking proxies. While it was not trained with any explicit reliability criterion, 22.0% of its 1,030 CpGs have high ICC (>0.75) versus only 8.5% of the array (log_2_ *OR* = 1.6, FDR = 8.5 × 10^−53^; Figure 4b), and poor ICC probes are significantly depleted (log_2_ *OR* = −1.56, FDR = 4.5 × 10^−69^). As previously shown by Sugden et al. (2020), probes with high test-retest reliability are fundamentally more likely to be consistently associated with a trait across EWAS studies. By pre-selecting probes based on their association with nine distinct traits, GrimAge effectively resolved this reliability bias. This finding suggests that pre-selecting CpGs based on association with virtually any strong biological trait will enrich for higher ICC probes, partly explaining GrimAge’s strong performance in external validation cohorts. Unsurprisingly, DunedinPACE, which explicitly excluded low-reliability probes during its construction, shows the most extreme depletion, with only 3.5% of its 173 CpGs having poor ICC (log_2_ *OR* = −5.76, FDR = 6.2 × 10^−66^).

### Concordance with Biological Literature

The atlas easily surfaces relationships that contextualize clock mechanics within the broader literature. For example, IntrinClock is significantly enriched for CoRSIVs (Correlated Regions of Systemic Interindividual Variation; log_2_ *OR* = 1.95, FDR = 1.7 × 10^−3^). This is a highly logical, albeit novel, finding: CoRSIVs are inherently defined by their correlated methylation status across different tissues within an individual, perfectly mirroring IntrinClock’s core design of filtering for probes that agree across varying cell types in blood. Similarly, RetroAge450K exhibits a distinct functional profile. It is the only clock significantly enriched in Open Seas (log_2_ *OR* = 2.61, FDR = 2.4 × 10^−211^) and Solo-WCGWs (log_2_ *OR* = 1.76, FDR = 2 × 10^−19^), while being significantly depleted for PRC2 binding sites as only 0.5% of its 1,317 CpGs overlap the top 10,000 PRC2-bound probes (log_2_ *OR* = −2.2, FDR = 1.8 × 10^−4^; Figure 4d). This perfectly mirrors the opposing dynamics of epigenetic aging, where PRC2-bound targets typically gain methylation over the lifespan while heterochromatic retroelements located in gene-poor open seas undergo progressive loss of methylation. Ultimately, the atlas proves its utility in verifying that specialized clocks are accurately capturing distinct mechanisms of biological aging.

### Novel Hypotheses for Further Investigation

Importantly, CpG Atlas quickly generates findings that warrant further investigation. For instance, the pace-of-aging clocks show striking enrichment for BMI-associated CpGs: 12.7% of DunedinPACE CpGs and 13.0% of DunedinPoAm38 CpGs appear in the EWAS Atlas BMI list (log_2_ *OR* = 4.56, FDR = 6 × 10^−86^ and log_2_ *OR* = 4.59, FDR = 9.2 × 10^−22^respectively; Figure 4c). However, BMI is also highly enriched in a specific, distinct subset of other clocks (Hannum, Zhang2019, GrimAge, and DNAmFI), raising an immediate biological question: what structural or training similarities cause these specific clocks to heavily capture metabolic risk compared to others? Furthermore, we observed that functional annotations such as histone modifications and transcription factor (TF) binding are enriched almost exclusively within chronological age clocks and PhenoAge. Understanding why chronologically trained algorithms disproportionately select for active regulatory regions, while mortality-trained clocks do not, remains an open avenue for future epigenetic research.

### Iterative biomarker discovery for inflammatory bowel disease using multi-layer filtering

Our second case study demonstrates how CpG Atlas supports systematic, quality-controlled biomarker discovery through sequential integration of multiple annotation layers. We constructed a four-step filtering workflow (Figure 5a) targeting methylation biomarkers for inflammatory bowel disease (IBD), including Crohn’s disease and ulcerative colitis. Each step applies a distinct biological or technical criterion from a different CpG Atlas layer, illustrating the kind of cross-layer integration that the CpG Atlas makes feasible.

**Figure 5.**
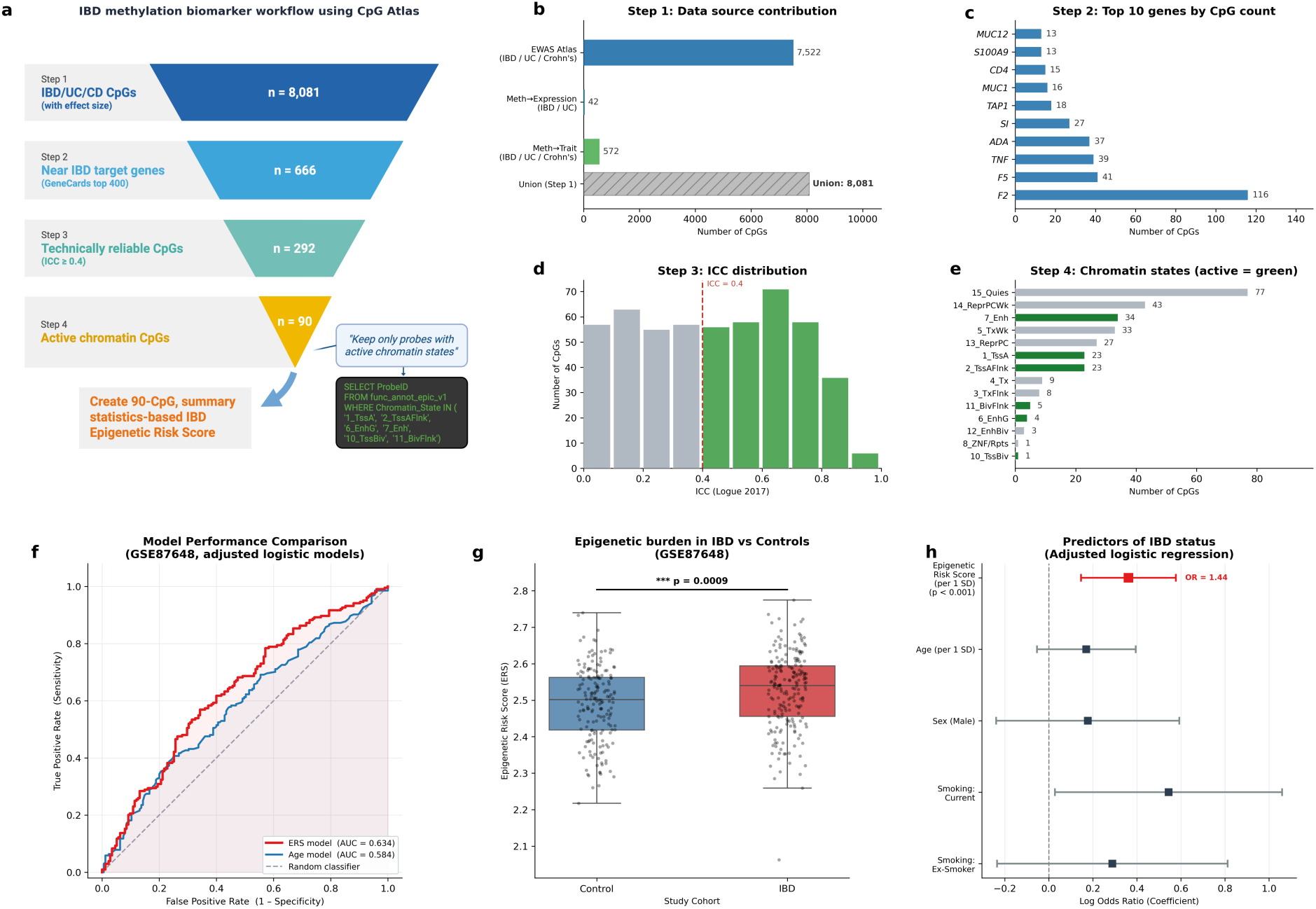
Iterative biomarker discovery. **a-e.** Four-step filtering pipeline applied to 8,081 IBD-associated CpG probes. Step 1: retain probes associated with IBD/Crohn’s/ Ulcerative Colitis (8,081 probes). Step 2: retain probes near IBD-relevant target genes (666 probes). Step 3: remove probes with low intraclass correlation coefficient (ICC < 0.4), reducing the set to 292 probes. Step 4: retain probes in active chromatin states (Enh, TssA, TssAFlnk, BivFlnk, EnhG, TssBiv), yielding a final set of 90 probes used to create a summary statistics-based IBD biomarker. **f.** Receiver Operating Characteristic (ROC) curves comparing the classification performance of the Epigenetic Risk Score (ERS) model (AUC = 0.634) against an age-only baseline model (AUC = 0.584) in an independent validation cohort (GSE87648). **g.** Unadjusted Epigenetic Risk Scores in GSE87648 healthy controls versus IBD patients (p < 0.001, ordinary least squares regression). **h.** Forest plot detailing the adjusted multivariable logistic regression model predicting IBD status. Continuous variables (ERS and Age) were standardized (per 1 SD) to allow for direct effect size comparison.

In Step 1, we queried three CpG Atlas IBD-related data layers: EWAS Atlas associations with traits matching IBD, Crohn’s disease, or ulcerative colitis (7,522 probes); EWAS Atlas methylation à gene expression causal links with IBD-related trait exposure (42 probes); and EWAS Atlas methylation à trait causal links with IBD-related outcomes (572 probes). The union of these three layers yielded 8,081 unique IBD/CD-associated CpGs with effect-size values available (Figure 5b). In Step 2, these CpGs were intersected with the EPICv1 manifest and filtered to probes annotated near the top 400 IBD-associated target genes identified via GeneCards relevance scores (49), reducing the set to 666 probes (Figure 5c). In Step 3, probes with ICC below 0.40 were removed, yielding 292 reliable probes (Figure 5d). In Step 4, the set was restricted to probes in active chromatin states (enhancers, active TSS, flanking active TSS, flanking bivalent TSS/Enh, genic enhancers, and bivalent TSS), producing a final set of 90 CpGs that satisfy all four criteria (Figure 5e; Supplementary Table S6).

The 90 final CpGs were used to construct an Epigenetic Risk Score (ERS) by extracting effect size weights from across all three IBD-related CpG Atlas data layers queried in Step 1: EWAS Atlas associations, methylation à gene expression causal links, and methylation à trait causal links, and computing the mean weight per probe across all contributing entries (Supplementary Table S6). The ERS for each sample was then calculated as the dot product of probe-level DNA methylation beta values and the corresponding mean effect size weights. To validate this score, we applied it to an independent dataset, GSE87648, comprising whole-blood DNA methylation profiles from IBD patients and healthy controls (50). ROC analysis yielded an AUC of 0.634, compared to a 0.584 AUC from the Age model (Figure 5f), reflecting modest but statistically meaningful discriminative capacity given that the score was completely derived from annotation-based filtering alone, without any sample-level training or cross-validation against this cohort. ERS values were significantly elevated in IBD cases relative to controls (Figure 5g), confirming that the multi-layer filtered probe set captures a meaningful epigenetic signal. To determine the independent predictive value of this signal, we evaluated the ERS in an adjusted logistic regression model incorporating age, sex, and smoking status as covariates. The ERS emerged as the only significant independent predictor of IBD status (OR = 1.44 per 1 standard deviation increase, p < 0.001; Figure 5h). In contrast, baseline clinical demographic variables did not demonstrate significant independent predictive value in this cohort, underscoring the distinct diagnostic utility of the methylation-derived risk score.

## Discussion

CpG Atlas addresses a significant limitation in DNA methylation research: the fragmentation of data and the absence of a unified, queryable framework for multi-dimensional interpretation of CpG sites. While several resources have made important contributions towards organizing DNA methylation knowledge, none integrates probe-level technical reliability, causal evidence, functional annotations, tissue specificity, repetitive element biology, and epigenetic clock membership within a single queryable database. The distinctive contribution of CpG Atlas is this centralization: when a researcher identifies a probe or list of probes of interest, answers to reliability, causal, functional, and clock questions are all retrievable from the same interface through the same query language, without any additional cross-database harmonization steps. This matters in DNA methylation research as these dimensions are not independent: a probe with high ICC, a causal association, active chromatin state, and membership in a mortality-trained epigenetic clocks is a very different candidate than a probe with a similar EWAS association but low reliability and open-sea location. Their joint interpretation, however, requires simultaneous access to all layers of CpG Atlas.

We present here two case studies to illustrate modes of analysis that CpG Atlas makes possible. The clock enrichment analysis demonstrates that cross-layer annotation reveals structural patterns in these clocks that are invisible from the clocks’ predictive accuracy results alone: while first-generation chronologiceven age clocks show no enrichment for highly reliable probes, clocks pre-selected for specific traits or trained on hard outcomes like mortality are systematically enriched for moderately to highly technically reliable probes and depleted for poor technical reliability probes, an emergent property of the training objective and not an explicit design criterion in many cases. Additionally, cross-layer analysis surfaces unexpected structural differences, such as the observation that chronologically trained algorithms are disproportionately enriched for active regulatory regions while mortality-trained clocks are not. The IBD biomarker workflow demonstrates the value of multi-step filtering for biomarker discovery: the 8,081-probe starting set shrinks to 90 candidates through a sequence of filters (gene proximity, reliability, chromatin accessibility) that each eliminate a distinct class of non-specific signal. The resulting Epigenetic Risk Score built from these 90 probes demonstrates significant, independent predictive value in an external patient cohort, providing robust biological validation. Both workflows reduce to a few SQL queries in CpG Atlas and would otherwise require hours of manual data harmonization across disconnected resources.

CpG Atlas addresses a fundamental challenge in applying large language models to molecular biology: the grounding problem. Standard LLMs process Illumina methylation probe IDs through sub-word tokenizers, meaning they have no intrinsic understanding of molecular identifiers. While a highly characterized probe (such as cg23124451) might possess strong statistical weight within the model due to frequent co-occurrence with specific genes or epigenetic clocks in the training corpus, the vast majority of the ∼850,000 probes suffer from highly sparse data representation. To a general-purpose model, these probes are mathematically indistinguishable from background noise. Consequently, when queried about these less-characterized CpG sites, LLMs suffer dangerously: rather than admitting ignorance, they generate highly confident, biologically coherent, yet entirely hallucinated functional annotations, especially if one of the sub-word tokens has a strong, random association with a specific gene or annotation in a totally different context. To illustrate this vulnerability, we queried four general-purpose LLMs about an obscure probe, cg00000321 (Table 1, full outputs available in Supplementary Data S1). The ungrounded models either provided very few relevant details with a generic, safe response (GPT-5.5, Gemini 3.1 Pro) or exhibited severe hallucinations, confidently misidentifying the probe’s chromosomal location, nearest gene, and regulatory context (Gemini 3 Flash, Claude Sonnet 4.6). However, when one of the worst-performing base models (Gemini 3 Flash) was augmented with retrieved tabular data from CpG Atlas, it achieved complete accuracy across all assessed metrics. Asking an LLM to rely on its parametric memory to reason about DNAm probes and their functional context is, therefore, highly flawed. A NL-to-SQL system on a deterministic database like CpG Atlas bypasses this vulnerability. By training on question-answer pairs grounded in the CpG Atlas schema, the model is no longer asked to retrieve biological facts from its highly uneven internal weights. Instead, it learns to interpret natural language queries in terms of specific tables, columns, and relationships that make up the database. This results in a system that achieves high accuracy not by memorizing answers, but by learning to translate the user’s intent into deterministic SQL queries. As the volume and complexity of epigenetic databases continue to grow, this approach to schema-aware fine-tuning provides a practical path toward AI systems that can genuinely reason about molecular data rather than merely pattern-match on its textual representation. More broadly, structured databases like CpG Atlas, where every claim is tied to a specific probe, study, and effect size, represent exactly the kind of deterministic knowledge architecture that future biomedical AI systems will require to produce reliable, verifiable, and hallucination-resistant outputs in the molecular domain.

**Table 1.** Comparison of 4 AI models in answering the question “What do you know about CpG site cg00000321? Give your answer in one paragraph.” Boxes are marked as “incorrect” if a model confidently stated a hallucinated annotation. Boxes are marked with a checkmark when a model answered correctly. Boxes are empty when a given annotation is missing in the AI’s answer.

|  | GPT-5.5 | Gemini 3.1 Pro | Gemini 3 Flash | Claude Sonnet 4.6 | Gemini 3 Flash + CpG Atlas |
| --- | --- | --- | --- | --- | --- |
| Chromosome | ✓ | ✓ | incorrect | incorrect | ✓ |
| Nearest gene |  |  | incorrect | incorrect | ✓ |
| Coordinates | ✓ | ✓ |  |  | ✓ |
| Relation to CpG Island | ✓ |  | incorrect |  | ✓ |
| Technical reliability |  |  |  |  | ✓ |
| Region |  |  |  |  | ✓ |
| Disease/Trait associations |  |  | incorrect |  | ✓ |
| Tissue specificity |  |  |  |  | ✓ |

We acknowledge several limitations in the current setup of CpG Atlas. First, coverage of array platforms remains uneven: ICC estimates, clock membership annotations, and EWAS associations are derived predominantly from 450K and EPICv1 studies, and EPICv2 and MSA probes have substantially less characterization in the published literature. Reliability estimates and enrichment analyses may not generalize across array generations due to differences in probe chemistry. Second, most of the annotation datasets we used are blood-derived, which limits the utility of CpG Atlas for researchers working in non-blood tissues such as brain or saliva. However, the process of adding information to CpG Atlas is straightforward; simply add a table to the SQL database, and therefore future results concerning other contexts, such as new arrays or other tissues, can be readily integrated into CpG Atlas. Third, while the NL-to-SQL interface makes database access much easier, the fine-tuned model was trained on a specific query distribution and might underperform on highly novel queries beyond its training scope, therefore we encourage users to inspect the generated SQL and make any necessary adjustments before treating results as definitive. Finally, the local deployment option requires sufficient computational resources to run a 7 billion parameter model, which may not be feasible in all research environments; users without compatible hardware are encouraged to use the API mode, which requires an external API key.

A key limitation of the current state of CpG Atlas is that the database reflects the current snapshot of the methylation annotation literature, and associations, reliability estimates, and functional annotations will inevitably evolve as new literature findings emerge. Addressing this, alongside expanding the preliminary hallucination assessment shown in Table 1, is a primary goal for future development. Specifically, we plan to systematically benchmark LLM performance rates across a highly diverse collection of CpG sites, quantifying the exact performance improvements and reduction in hallucination by CpG Atlas integration. Planned work also includes a versioned release pipeline that tracks changes in schema and data inventory across updates, automated monitoring of preprint and publication repositories for new EWAS, clock, ICC studies, etc. In the near term, users can already extend the database with their own annotation layers by uploading a CSV or XLSX file with a ProbeID column and a structured data description, enabling integration of unpublished or lab-specific data without requiring database expertise. More broadly, we envision CpG Atlas as a living infrastructure resource rather than a fixed release, a database that grows with the field and provides the kind of up-to-date, structured biological grounding that modern AI-assisted research workflows increasingly require.

## Methods

### Database construction and data curation

CpG Atlas is built on DuckDB (27), an open-source, in-process analytical database management system for fast Online Analytical Processing (OLAP). The database comprises 30 harmonized tables organized across six functional categories. The complete database covers over 1.2 million unique CpG sites across the HumanMethylation450, MethylationEPIC v1.0, MethylationEPIC v2.0, and the Methylation Screening Array platforms. Array manifests were obtained directly from Illumina. Each of the remaining annotation layers was sourced from its primary publication and processed to conform to a common schema using a standardized Python pipeline with probe identifier as the primary key. Cross-platform mapping information from array manifests was used to harmonize probe identifiers across array generations.

### CpG Atlas Explorer web interface

The CpG Atlas Explorer is implemented using a single self-contained HTML application leveraging DuckDB-WebAssembly (DuckDB-WASM) (46) to execute SQL queries client-side without server infrastructure. Database tables are exported to Apache Parquet format and hosted on Cloudflare R2 object storage. The explorer is deployed via Vercel at *cpg-atlas-pi.vercel.app*. Users may upload their own tabular data (CSV or Excel) and join it against any Atlas table via the SQL Query module.

### Natural language query interface

The fine-tuned NL-to-SQL system is based on Qwen2.5-Coder-7B-Instruct (47) fine-tuned with Low-Rank Adaptation (LoRA) (48) with 4-bit quantization on a curated dataset that merges CpG Atlas-specific question-answer pairs with dynamic subsets of the database schema documentation, formatted in ChatML so the model learns to read and apply schema context at inference time. Training converged from an initial loss of 1.62 to 0.005, confirming successful schema internalization. The fine-tuned weights are publicly released to support local deployment. The system can operate in two modes: Local Mode (fine-tuned Qwen runs on user hardware, no API key required) and API Mode (use-supplied key for OpenAI, Anthropic Claude, or Google Gemini). To test the accuracy of different models in translating CpG Atlas natural language queries into SQL, we used LLM-as-a-Judge to evaluate model outputs on a held-out set of question-answer pairs and classify them as correct, partially correct, or incorrect. We ran this evaluation on the base Qwen2.5-Coder-7B, the fine-tuned Base Qwen2.5-Coder-7B, and GPT-5.4 mini. LLM-as-a-Judge correctness evaluates whether the generated SQL and result fully answer the question, independent of the ground truth SQL. Partially correct scores are given to near-miss outputs, such as queries with minor repairable schema-name errors or queries that answer only part of a multi-condition question. Because partial credit may benefit less precise models, it is treated as a secondary diagnostic metric rather than the primary accuracy endpoint. Generated SQL queries that do not answer the question or do not execute (due to errors other than table/column naming) are graded as incorrect.

### Epigenetic clock enrichment analysis

Enrichment was assessed using a Yates-corrected chi-squared approximation of Fisher’s exact test, with Benjamini-Hochberg FDR correction applied across all 336 tests (16 clocks x 21 annotations). The statistical background comprised all valid assay probes on the 450K array (N = 485,512). The Yates continuity correction was utilized to ensure conservative P-value estimates given the large background size, with exact probabilities approximated via the Abramowitz and Stegun polynomial expansion of the complementary error function. Effect sizes are reported as the base-2 logarithm of the Odds Ratio (log_2_(*OR*)). Cells with zero clock-annotation overlap with an undefined OR are plotted as hatched and were excluded from multiple testing correction. Clock CpG lists were derived from the methylCIPHER R package (25). The 16 clocks selected span chronological age (Hannum (12), Horvath1 (11), Horvath2 (51), Zhang2019 (52)), telomere length proxy (DNAmTL (53)), biological age and mortality (PhenoAge (13), GrimAgeV1 (14)), pace of aging (DunedinPoAm38 (54), DunedinPACE (18)), causal aging (CausAge, AdaptAge, DamAge (19)), immune composition (DNAmIC (55)), frailty index (DNAmFI_Li (56)), cell-intrinsic aging (IntrinClock (40)), and retroelement-based clock (RetroAge450K (20)). Full enrichment statistics for all 336 clock–annotation pairs are provided in Supplementary Table S4.

### IBD methylation biomarker discovery workflow

To identify robust methylation biomarkers for Inflammatory Bowel Disease (IBD) and Crohn’s Disease, candidate CpG sites were processed through a four-step filtering pipeline using CpG Atlas. First, candidate CpGs associated with IBD, Crohn’s Disease, or Ulcerative Colitis were aggregated from three EWAS Atlas tables: direct trait associations, causal methylation-to-expression, and causal methylation-to-trait models. Only probes with non-missing effect sizes were retained, yielding an initial pool of 8,081 CpGs. Second, to ensure biological relevance, this pool was restricted to 666 CpGs mapped to the top 400 IBD-associated target genes (from GeneCards) using the EPICv1 array manifest. Third, to optimize technical reliability, probes were filtered based on their Intraclass Correlation Coefficient (ICC) from the Logue et al. (2017) study. Probes with an ICC ≥ 0.4 were retained, reducing the set to 292 CpGs; probes with no ICC data in the reference set were also retained. Fourth, we selected for functional genomic context by restricting the panel to CpGs located within active chromatin states (Enh, TssA, TssAFlnk, BivFlnk, EnhG, TssBiv), yielding a final, highly specific set of 90 biomarker probes. An Epigenetic Risk Score (ERS) model was constructed using this final subset of 90 CpGs. For each CpG, the initial effect sizes retrieved from the EWAS Atlas sources in Step 1 were compiled. For probes appearing in multiple sources, model weight was calculated by taking the mean of the available effect sizes. The ERS was validated using an independent patient cohort (GEO accession GSE87648). Prior to ERS calculation, missing DNA methylation beta values in the validation cohort were mean-imputed at the individual probe level. A patient-specific ERS was then calculated as the dot product of the individual’s methylation profile and the probe weights. Statistical evaluations of the ERS were conducted using regression frameworks. First, the classification performance of the IBD ERS was quantified using Receiver Operating Characteristic (ROC) curves, comparing the Area Under the Curve (AUC) of a full adjusted ERS model against a baseline clinical model (Age, Sex, and Smoking status only). Next, differences in the unadjusted ERS between IBD cases and healthy controls were assessed using Ordinary Least Squares (OLS) regression. Finally, to evaluate the independent predictive value of the ERS, a multivariable logistic regression model was fitted to predict IBD status. This model incorporated the ERS alongside relevant clinical covariates, including sex and smoking status (current and ex-smoker). To facilitate the direct comparison of effect sizes within this model, both continuous variables (patient age and ERS) were Z-score standardized (per 1 standard deviation) prior to model fitting.

## Data and code availability

The CpG Atlas database tables are publicly available through the CpG Atlas Explorer webtool accessible at *<u>cpg-atlas-pi.vercel.app</u>.* LoRA adapter fine-tuned on Qwen2.5-Coder-7B-Instruct is available at *<u>huggingface.co/vandijklab/CpGAtlas-NL-to-SQL-Qwen2.5-Coder-7B-LoRA</u>.* The full .db file and code used to build it will be available upon publication.

## Acknowledgements

This work is supported by the National Institute on Aging (NIA:2R01AG065403 to A.H.C.) and National Institutes of Health (NIH) grant (R35GM143072-01 to D.v.D.). This work is also supported by an AI at Yale Seed Grant to A.H.C. and D.V.D. J.F.A. is additionally supported by the Biomarker Network Pilot Project Award (R24 AG037898) and the Gruber Science Fellowship at Yale University. N.E. is supported by a National Health and Medical Research Council (NHMRC) Investigator Grant (APP1194159), an ARC Discovery Project grant (DP240102155), and a Hevolution/AFAR New Investigator Award in Aging Biology and Geroscience Research. The authors would like to thank the researchers who made their study data publicly available and acknowledge that without their data, this project would not have been possible. BioRender (https://BioRender.com) was used for Figures 2 and 5.

## Author Contributions

J.F.A. and A.H.C conceived the project and study design. J.F.A built the CpG Atlas database and webtool and performed case-study analyses. S.W. and J.F.A. designed the NL-to-SQL architecture. S.W. fine-tuned the Qwen model. M.J. and N.E. provided Tissue-Specific Aging DMP module data. D.B., R.S, S.R., H.Z., and D.V.D. assisted with study design and contributed intellectually through manuscript editing and reviewing findings. J.F.A. and A.H.C. wrote the manuscript, and all authors reviewed and contributed to the manuscript.

