## Supplementary_Data for "CpG Atlas: A centralized multi-layer database and AI interface for DNA methylation research": Supplementary_Data_S1.docx

Prompt: What do you know about CpG site cg00000321? Give your answer in one paragraph.

GPT-5.5: CpG site cg00000321 is an Illumina methylation-array probe targeting a CpG dinucleotide on human **chromosome 8** at approximately **hg19 position chr8:41,167,802**, located in a **CpG “shore” region** rather than within a dense CpG island. Current public annotation resources **do not assign it to a well-characterized nearby gene**, and EWAS Atlas presently lists no strongly established trait associations for this probe. Like other CpG loci, methylation at cg00000321 can potentially vary across tissues and biological states, and available datasets show measurements from many tissues including blood, brain regions, adipose tissue, liver, placenta, sperm, and immune cell types. In epigenetics studies, probes such as cg00000321 are commonly analyzed for differential DNA methylation linked to aging, disease, environmental exposure, or cell-type composition, although this specific site does not appear to be among the best-known disease-associated CpGs in the literature.

Gemini 3.1 Pro: CpG site cg00000321 is a specific DNA methylation probe identifier standardly utilized in Illumina Infinium arrays, such as the HumanMethylation450 and 850K EPIC platforms. Mapped to human **chromosome 8** (specifically at **hg19 coordinate chr8:41167802**), the methylation status of this locus is commonly quantified as a beta value in epigenome-wide association studies (EWAS). Structurally, it is situated in extremely close proximity to a known single nucleotide polymorphism (**SNP rs144697833**, located just two base pairs away), a variable that researchers often account for during quality control to prevent probe-binding artifacts from skewing methylation readouts. Furthermore, while many highly predictive CpGs feature dense regulatory architectures, recent motif mapping analyses have cited cg00000321 as an **example of a methylation site that lacks recurrent transcription factor binding motifs** (such as ARNT or Forkhead), suggesting that its variation in methylation may lack the direct, multi-tissue regulatory impact characteristic of classical promoter-bound epigenetic markers.

Gemini 3 Flash: cg00000321 is a specific CpG site targeted by the Illumina Infinium HumanMethylation arrays (such as the 450k and MethylationEPIC beads) and is located on **chromosome 1** within the **GPR153 (G Protein-Coupled Receptor 153) gene**. Positioned in a **CpG island** within the gene's **promoter region**, its methylation status is frequently scrutinized in epigenome-wide association studies (EWAS) due to its potential role in gene silencing and regulatory control. Research has often linked the methylation levels of this site to various biological processes and conditions, including **neurodevelopmental traits**, **aging**, and certain **metabolic responses**. In bioinformatics workflows, it is commonly analyzed using tools like minfi or limma to determine differential methylation patterns across diverse tissue types or clinical cohorts.

Claude Sonnet 4.6: The probe identifier cg00000321 follows the naming convention used by Illumina's Infinium methylation array platforms (such as the HumanMethylation27K, 450K, and EPIC arrays), where each "cg" prefix designates a CpG dinucleotide site interrogated by a specific probe on the array. According to the Illumina manifest files and associated annotation resources (such as those hosted at the UCSC Genome Browser or the MethylationEPIC manifest), cg00000321 maps to **chromosome Y**, within or near the **gene TSPY4 (Testis Specific Protein Y-Linked 4)**, and is measured as a beta value representing the proportion of methylated alleles at that cytosine position in a given sample. Because it resides on the Y chromosome, this site is only meaningfully assayed in male samples, and it is typically used in sex-check quality control steps during array-based DNA methylation analyses. It has appeared in various epigenome-wide association study (EWAS) pipelines, though it is generally excluded from autosomal analyses or flagged due to its Y-linkage and the associated cross-hybridization concerns that affect sex chromosome probes on these arrays.

Prompt: Given attached tables, what do you know about CpG site cg00000321? Give your answer in one paragraph.

Gemini 3 Flash: The CpG site cg00000321 is an **Infinium Design Type II** probe located on **chromosome 8 (41,310,282 in hg38)**, associated with the **SFRP1 gene** within the **TSS1500 region** and situated in a **South Shore (S_Shore)** relative to a CpG island. It demonstrates variable but generally high reliability across platforms, with Intraclass Correlation Coefficient **(ICC) values ranging from 0.48 to 0.79**, including an "Excellent" reliability rating for the EPICv1 array. This site is a significant biomarker for several conditions, showing strong associations with **Alzheimer's disease** (p=2.35×10^-6), **B Acute Lymphoblastic Leukemia**, and **meningioma deterioration**. Furthermore, it acts as a tissue-specific hypermethylated differentially methylated probe (hyperDMP), showing highly significant positive methylation across various tissues, most notably in **Blood** (p=1.6×10^-30), as well as in **Skin**, **Liver**, and **Colon**.
